# Temporal integration of rod signals in luminance and chromatic pathways

**DOI:** 10.1101/2022.03.17.484829

**Authors:** Iñaki Cormenzana Méndez, Andrés Martín, Beatriz O’Donell, Dingcai Cao, Pablo A. Barrionuevo

## Abstract

We assessed how rod excitation (R) affects luminance (L+M+S) and chromatic [L/(L+M)] reaction times (RTs). A four-primary display based on the overlapped images of two spectrally-modified monitors, which allowed specific or combined [L+M+S+R, L/(L+M) + R] photoreceptor stimulation, was used to present a C-target stimulus differing from the background only by the selected stimulation. For the luminance pathway, rod input increased RTs, suggesting a suppressive rod-cone interaction. The responses of the chromatic pathway were faster when rods were involved, suggesting a major role of rods in mesopic color perception.

## 1. Introduction

Retinal information is conveyed to the lateral geniculate nucleus in the visual stream via postreceptoral pathways; the magnocellular (MC) and the parvocellular (PC) are two major retinogeniculate pathways. The MC-pathway begins in the retina with the diffuse bipolar cells and the parasol ganglion cells; it codifies the luminance information of the visual scene conveying additive L-, M- and S-cone signals [1]. Instead, the PC-pathway codifies chromatic opponency (roughly *red-green*) through antagonist L- and M-cone signals, followed by processing in the midget bipolar and ganglion cells. It was demonstrated that the MC-pathway responds predominantly to temporal frequencies as a band-pass filter, rather, the PC-pathway has a low-pass filtering characteristic [2,3]. In mesopic and scotopic light levels, rods contribute to visual perception. In primates, rod signals do not preserve a private pathway to reach other brain centers outside the retina, instead, they share postreceptoral circuits with cone signals. Rod signals are thought to be conveyed by two pathways: one pathway via ON rod bipolar cells, AII amacrine cells, and ON and OFF cone bipolar cells; and the other via rod-cone gap junctions (coupling) and ON and OFF cone bipolar cells. The role of these two pathways is thought to vary with light level and the gap junction pathway is faster than the AII pathway [4–6]. Rod signals are slower than cone signals; this delay is dependent on the rod pathway involved. For the AII pathway, the delay could be up to 80 ms [7,8], while for the gap junction pathway the delay is much lower [9] and it was estimated in the range of 8 - 20 ms [10,11].

Physiological studies showed that the MC-pathway predominantly conveys rod signals [12,13]. There is also evidence of rod intrusion to midget cells of the PC-pathway [13,14]. Recent studies using multi-primaries photostimulators have contributed to a better understanding of how rod and cone signals combine in specific visual pathways (for a review see [6]). These systems allow modulating independently the excitation of selected photoreceptors through the method of silent-substitution [15,16]. A major advantage of these systems is the evaluation of independent photoreceptor contribution at the same adaptation level. Using this type of system, it was found that rods and cones are combined in a vector summation way in the MC-pathway [11,12], and through probability summation in the PC-pathway [11]. Rods intrusion produced a hue percept shift to green and blue [17], and cone chromaticity discrimination is evidently deteriorated with rod intrusion [18], which is further evidence of rod intrusion to the chromatic pathways. Also, it has been suggested that lateral rod-cone interaction increased cone reaction times [19]. Therefore, in mesopic conditions rod intrusion should affect temporal dynamics of MC- and PC-pathways. Furthermore, rod influence could be different for each pathway. However, the nature of rod intrusion in the PC-pathway was not clearly established as it was for the MC-pathway, and it remains one of the main questions in mesopic vision [20].

Also, it is not clear the temporal dynamics with respect to the ON and OFF retinal processing. It has been suggested that ON signals are slower than OFF signals in the peripheral retina [2,10,11,18,19,21–23]. However, there is also opposed evidence [24,25]. Therefore, extrafoveal reaction times might contribute to clarify this behavior.

Due to the sluggish nature of rod signals, a mesopic vision complexity appears in field tasks where transient situations challenge the visual system [5]. To understand how visual perception, in particular luminance and color, is affected by rod intrusion in real-world mesopic conditions, a reliable approach is to study reaction time measurements. The objective of this study was to determine the contribution of rods signals in the temporal processing of the MC- (luminance) and PC- (chromatic) pathways. Therefore, we assessed rod-cone interactions in postreceptoral pathways using reaction times (RTs).

## 2. Materials and methods

### Observers

Three observers with normal or corrected to normal visual acuity participated in the experiment. One experienced psychophysical observer (male, 34 years old) and two naïve observers (2 females, 32 and 27 years old). Observers were monocularly tested. The observers had a normal color vision as evaluated by the Nagel anomaloscope. The study protocols were approved by the Institutional Review Board at the University of Illinois at Chicago and were in compliance with the Declaration of Helsinki. We obtained written consent for all observers. All of the procedures were carried out in accordance with US government human observer protection guidelines and regulations.

### Apparatus

The Maxwellian-view four-primary photostimulator introduced for studying rod-cone interactions [11,26] has been a major technical contribution to the visual field [27]. Similar approaches added a fifth primary in order to isolate the contribution of melanopsin activation and to study its integration with rods and cones [28,29]. However, these systems have a limited spatial resolution. To overcome this limitation, new approaches have recently emerged to apply silent-substitution in screens in which each pixel can be treated as a multi-primary light source [30–33]. However, these new approaches were not used to test visual performance in the mesopic region so far.

A calibrated CRT-based four-primary photostimulator controlled the excitations of rods and three types of cones (S-, M-, and L-cones) independently using the principle of silent substitution as defined by [16]. Effectively, the four-primary wavelengths and intensities are carefully chosen to maintain the excitation levels of some photoreceptor classes while varying the excitation of specific photoreceptor classes using silent substitution [15]. CRTs monitors were used to allow independent control of rods and cones excitations in each pixel in the display, so each pixel was a four-primary based light source [31]. To achieve this, we superimposed two CRT monitors through a partially silvered glass (see Fig. 1A for the optical layout). The two CRTs were covered by different cut-off spectral filters in order to obtain primaries with different spectral distributions. We used blue (B) and green (G) phosphors from the first CRT and green (G) and red (R) from the second CRT (see Fig. 1B, first row, for their spectral distributions). To produce two G primaries with different dominant wavelengths for both CRT displays, spectral cut-on, and cut-off filters were used, “R361Blue” (Rosco Laboratories Inc.) that cuts off wavelength between 550–650 nm wavelengths for CRT1 and “R16Amber” (Rosco Laboratories Inc.) that cuts off 450–520nm wavelengths for CRT2. (Fig. 1B, second row, for their spectral transmittance characteristics). The results are G1 for CRT 1 with a dominant wavelength of 518 nm and G2 for CRT 2 with a dominant wavelength of 558 nm (Fig. 1B, third row).

**Fig. 1.**
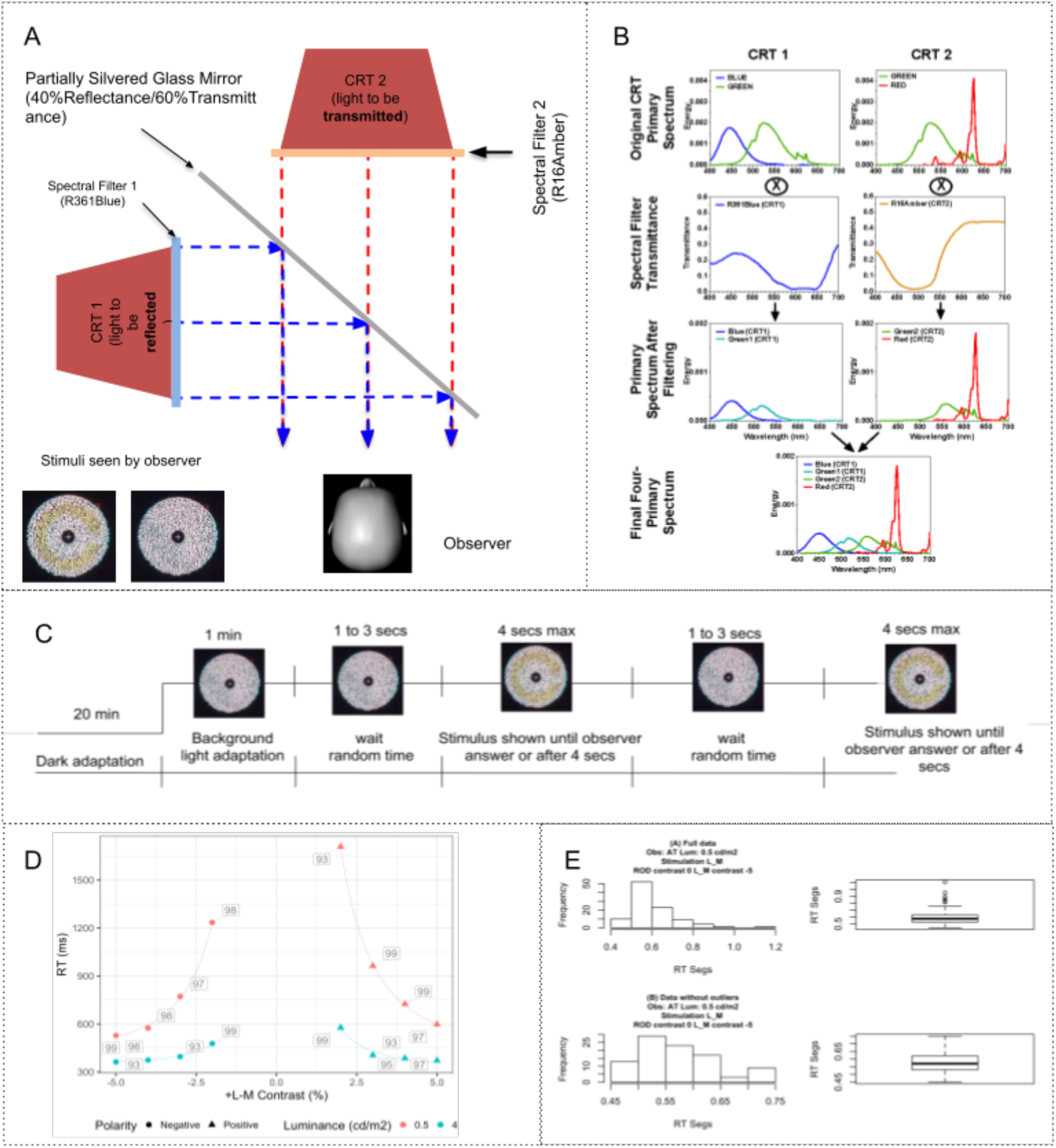
Experimental setup. (A) CRT-based four-primary photostimulator optical layout with stimulus example. (B) Primary Spectrum for the two CRTs and the filters. (C) Temporal sequence of the stimulus presentation. (D) Example of reaction time of +L-M cones condition for an observer. Percentage of correct answers are shown for each result. (E) Example of histogram for one combination before (upper row) and after (lower row) cleaning outliers with a cutoff criterion of 1.96 of standard deviation.

The same image is presented in the two CRT displays, so the image in CRT 1 is displayed using B and G1, and the image in CRT 2 is displayed using G2 and R. Then the two images are superimposed using a Teleprompter Glass mirror (40% reflectance for CRT1 and 60% transmittance for CRT2, Fig. 1A). The images are perfectly aligned at each corresponding pixel. This results in a design of four primaries (B, G1, G2, R) at each pixel (Fig. 1B, bottom row).

To control three displays simultaneously, including the two CRTs and one main display, we used a Macintosh Pro Desktop (Apple Inc.) that hosts an ATI Radeon HD 5770 video card (Advanced Micro Devices, Inc). Computer software has been developed in Matlab R2016a (Mathworks Inc.) with Psychophysics Toolbox Version 3 (PTB-3) and to allow individual control of each primary at each pixel. A two-button joystick latency was measured resulting in a meantime latency of 8 ± 3 ms, for both buttons.

Visual stimuli generated using a four-primary photostimulator were specified with L-cone, M-cone, S-cone, and rod (R) excitations [16,34]. We normalized rod excitations such as for a light metameric to an equal energy spectrum (EES), 1 photopic Troland (Td) is equal to 1 rod Td. The cone chromaticity is described in a relative cone-Td space, which plots *l* = *L*/(*L*+*M*) versus *s* = *S*/(*L*+*M*) with a normalization so that *l* = 0.667 and *s* = 1.0 for an EES light [35].

The device was fully calibrated considering spectral distribution, linearity, and spatial homogeneity to ensure accurate presentation of visual stimuli. After physical calibration, we conducted an observer calibration in order to compensate for the individual differences in photoreceptor spectral sensitivity and lens transmittance using the method of heterochromatic flicker photometry (HFP) at 15 Hz. We conducted additional measurements to confirm the photoreceptor isolation using a spectroradiometer PR-670 (PhotoResearch Inc.) to verify the correct computation in each stimulus condition [36].

### Stimuli

The stimuli were a 12° circular field with a “C” target, subtending 2° in the gap, 9.6° in the outer diameter, and 6.4° in the inner diameter; therefore, the mean radial eccentricity was 4°. A fixation cross was located at the center of the field at a viewing distance of 88 cm. The “C” target differed from the field only by its selected stimulation along with 1) the luminance (L+M+S), we added S to L+M in order to maintain an exact isochromatic percept [12], this condition codifies the inputs to the MC-pathway [1]; 2) the cone combination *L*/(*L* + *M*) which represents variation in the horizontal axis of the MacLeod and Boynton cone chromaticity space [35] and codifies variation in the PC-pathway [1]. The “C” target and background were made up of five types of discrete circles (0.48°, 0.28°, 0.52°, 0.36°, 0.32°) with the same luminance. The fixation cross also had the same luminance and stimulation as the background.

Experiments for the MC-inferred pathway were conducted at four mesopic luminances. Two in the high mesopic range (6 cd/m2 and 4 cd/m2), and two at the intermediate mesopic range (0.75 cd/m^2^ and 0.5 cd/m^2^), which were obtained with a 0.9 log unit neutral-density (ND) filter placed between the eye and the stimulus. Measurements with a 1.8 log ND filter, could not be perceived by the observers. An analysis for the difference between 6 cd/m^2^ and 4 cd/m^2^ didn’t show any effect of this difference. Therefore, we group these two luminances and we call it “High light level”. Similar results were found for 0.75 cd/m^2^ and 0.5 cd/m^2^ and we call this group “Low light level”. For the PC-inferred pathway the “High light level” was 4 cd/m2 and the “Low light level” was 0.5 cd/m2.

Depending on the stimulus conditions, we measured RTs for different Weber contrasts for both visual pathways (Experiment 1 and 2). For the MC-inferred pathway, in the interaction, the three conditions (L+M+S, R, and L+M+S+R) were stimulated with the same contrasts (−21% to 27%) (Table 1). For the PC-inferred pathway, the experiment was carried out under different contrasts for cones (±5% to ±2%) than rods (±25% to ±15%) in the three conditions (L/(L+M), L/(L+M)+R, R) due to gamut limitations and due to the different nature of the activation threshold (Table 1). For the same reason, the background chromaticity is different for positive and negative contrasts for this inferred pathway.

**Table 1.**
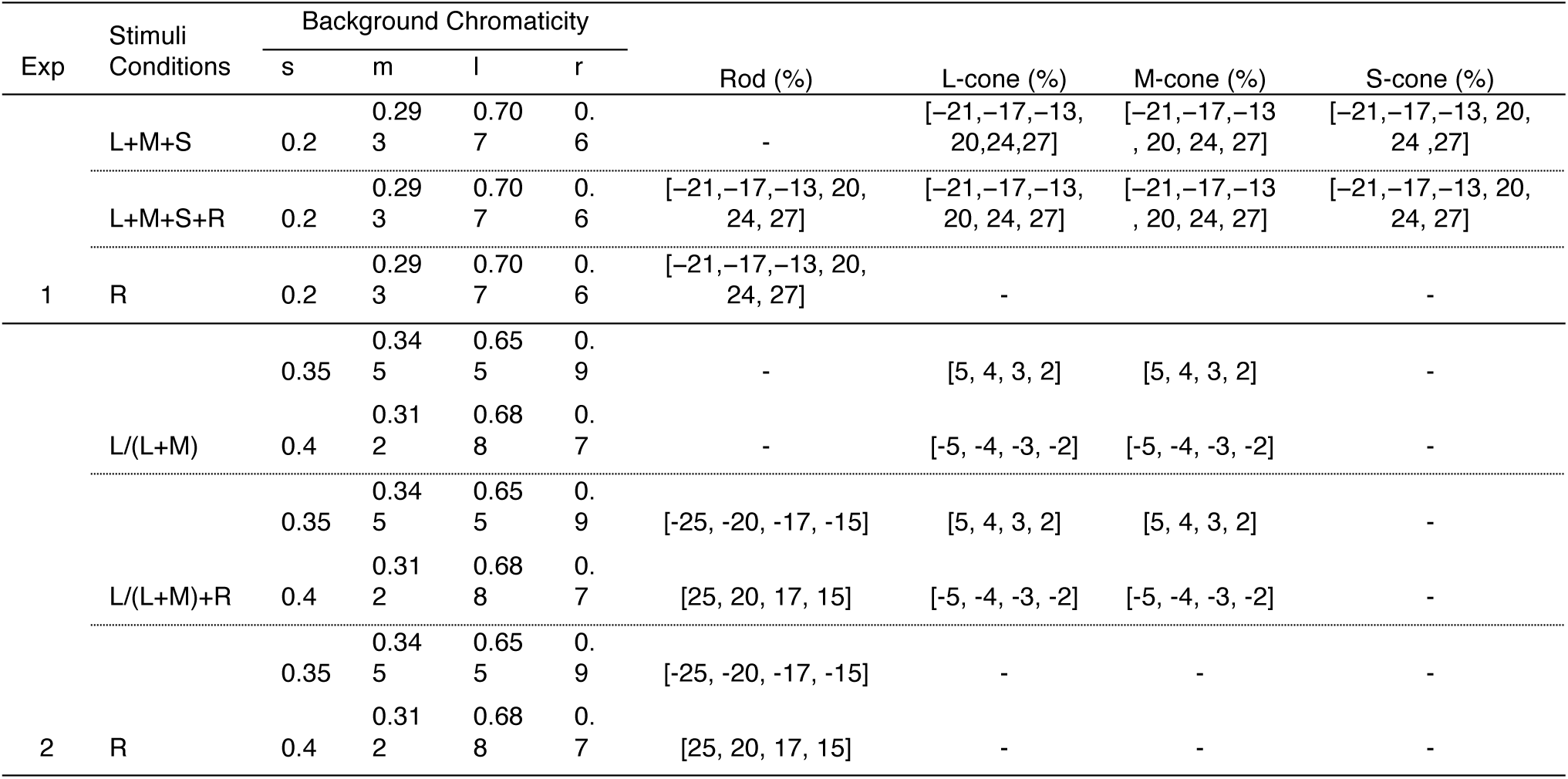
Stimuli Conditions for both experiments. The values of the relative photoreceptor excitations are expressed as a percentage of Weber contrast (%).

### General Procedure

The experiments were conducted in a dark room. During each trial, the observers were asked to fixate on the cross in the center of the stimuli (Fig. 1). Observers were monocularly tested and the head position was maintained using a chin rest. All testing sessions began with an initial 20 min dark-adaptation period and followed by an experimental session. The temporal procedure is shown in Fig. 1.

A “C” target stimulus presentation was preceded by a 1 min. light adaptation period to background chromaticity and a random time interval that varied between 1000 and 3000 ms, to avoid expectancy. Then, the “C” target was displayed and this triggered a timer to start until the participant answered. A two-alternative forced-choice task was used. The “C” gap was shown in two alternate orientations -at the right or to the left of the fixation cross-. To measure the RT, the participant pressed a two-button joystick to indicate the “C” gap orientation (right or left) as soon as he/she had detected the “C” target position. If the observer does not respond after 4 seconds, the C target disappears but the background was displayed. We discard these responses from the analysis for now. The time between the presentation of the stimulus and the observer’s response was registered as the RT. The “C” target stimulus was turned off when the observer’s response was recorded but the background was maintained. Once the observer responded, the 1000–3000 ms random time interval was reintroduced, and, during this period, the mean luminance was kept constant. Each experimental session took around one hour.

In a single session, a stimulus condition was measured at multiple contrasts at one light level. The stimulus conditions were randomized within and across sessions. A single run consisted of 36 combinations (2 light levels *6 contrasts * 3 stimulations = 36 conditions) for MC-inferred pathway, and of 96 combinations {2 light level*[(4 contrasts*2 polarities*2 stimulations) + (4 cone contrasts * 4 rod contrasts*2 polarities)]} for PC-inferred pathway.

A minimum of 100 repeats per combination was recorded. If the observer correctly answers the position of the C-target gap, it is considered a correct answer.

### Data Analysis

Reaction Time (RT) data constitutes a complex set of numbers that require, in order to avoid some known biases, a two-step process: 1) cleaning and inclusion criteria; 2) modeling and fitting.

Inclusion criteria: at each stimuli intensity level, only those intensities that triggered hit rates above 95% were considered. It was also assumed a normal distribution of the RT data for each stimulus intensity. Thus, a 1.96 standard deviation metric was applied as an outlier criterion: those RTs above or below 1.96 SD from the mean were discarded.

Modeling: raw RT data is modeled through the nonlinear Pieron equation. An exponent equal to 2 is assumed, this value was recommended for isoluminant stimuli [37,38]. The Pieron equation that we will use in the work takes the following form:

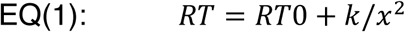

Where RT is the response time measured in the perceptual task. RT0 encodes the base reaction time: the asymptotic value to which the function tends when the signal intensity increases. k is the gain and x represents the signal intensity [39], i.e. Weber contrast (see Stimuli).

Taking 1/x^2^ as the value of the signal it is possible to linearize the ratio. In this way, the final equation is

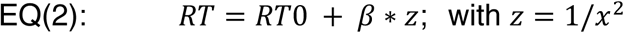

One advantage of the linear form of the relationship between reaction time and signal intensity is that the analysis and interpretation of the data are simplified since it allows applying linear models (e. g. ANCOVA). Within the linear model framework, the intercept encodes the RT0; the slope is the inverse of the gain: every increment in the X-axis implies a decrement in the stimuli contrast thus, an increment in the difficulty to perceive the orientation of the C and, therefore, an elevation of the reaction time.

Linear relationships also allow us to define comparison criteria. For example, a non-symmetric response of the MC or PC inferred channel can be assessed when there is a significant difference for positive respect to negative contrast conditions between slopes. *RT*_0_ is primarily determined by the time of the motor response, initiated when the sensory component reaches a criterion value. The parameter *RT*_0_ incorporates fixed components such as synaptic delays and conduction times [40]. A similar approach was implemented for assessing the effect of the light level. We applied this linear analysis to our results in the different conditions tested.

## 3. RESULTS

### Experiment 1: Intrusion of rods in the MC-inferred pathway

The contributions of rods, in terms of reaction times RTs, to the MC-inferred pathway (orange dots), for observers Obs. 1 (Panel A) and Obs. 2 (Panel B), are shown in Fig. 2. Also, results for the isolated MC-inferred pathway (*L+M+S*, green dots) and isolated rods (*R*, purple dots) conditions are shown in Fig. 2. Overall for each stimulation condition, RTs decreased with excitation contrast increments, as was expected [38].

**Fig. 2.**
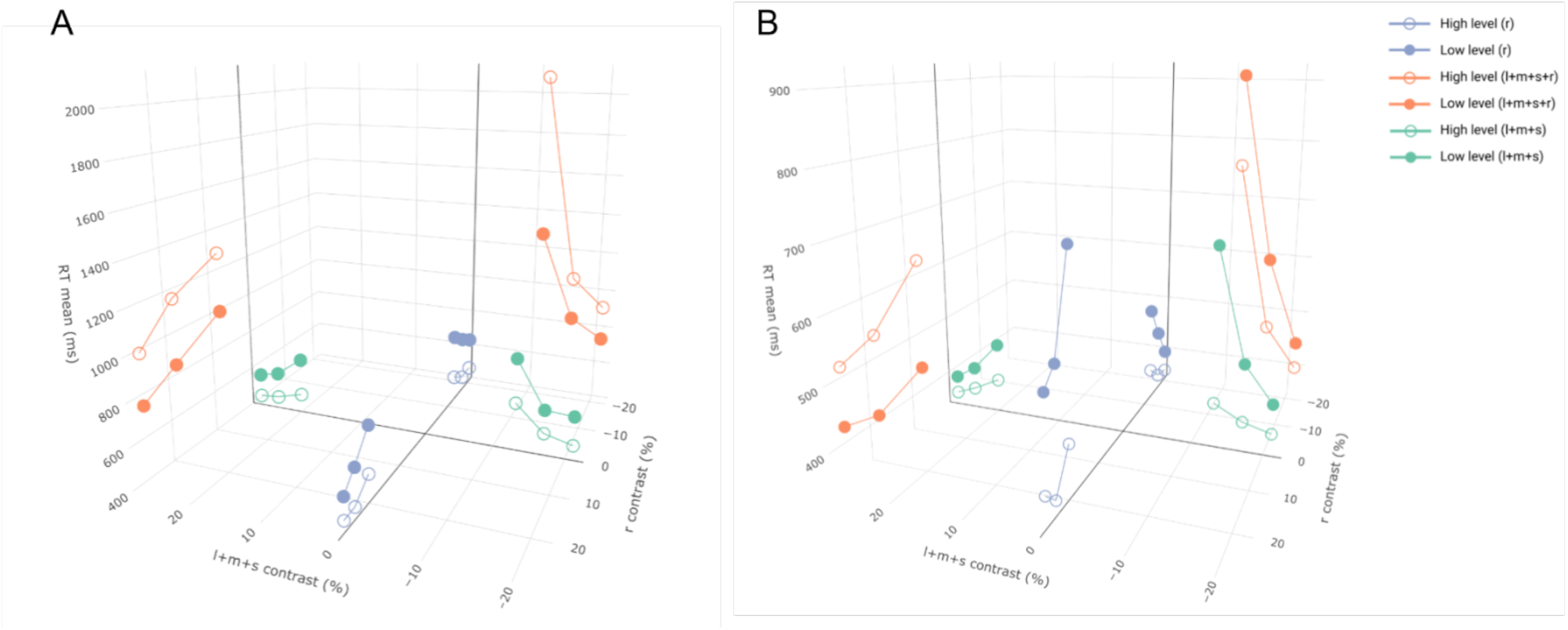
Three-dimensional graph showing the reaction times for the MC-pathway with the independent variables of excitation contrast of cones (*L+M+S)* in a horizontal axis and rods (*R*) on the other horizontal axis. Data is represented in the Pieron’s space (before linearization). The vertical axis represents the mean of the reaction times in milliseconds [RT mean (ms)]. Results are shown for isolated rod condition (purple dots), isolated MC-inferred pathway condition (green dots), and intrusion of rods to MC-inferred pathway condition (orange dots).

#### Isolated contributions (L+M+S, R)

##### Polarity effect

For the L+M+S condition (Fig. 3, upper panels), no differences were found for the intercept (Obs. 1: *t* = *0*.*11, p* = 0.925; Obs. 2: *t* = −1.27, *p* = 0.332) nor for the slopes (Obs. 1: *t* = −0.70, *p* = 0.557; Obs. 2: *t* = 0.41, *p* = 0.724) at low light level. Instead, for the high light level significant differences were found for both parameters from observer 2 (intercept: *t* =14.82, *p* < 0.01; slope: *t* = −19.64, *p* < 0.01) but no differences were found for observer Obs. 1 (intercept: *t* = −1.04, *p* = 0.409; slope: *t* = 0.41, *p* = 0.720).

**Fig. 3.**
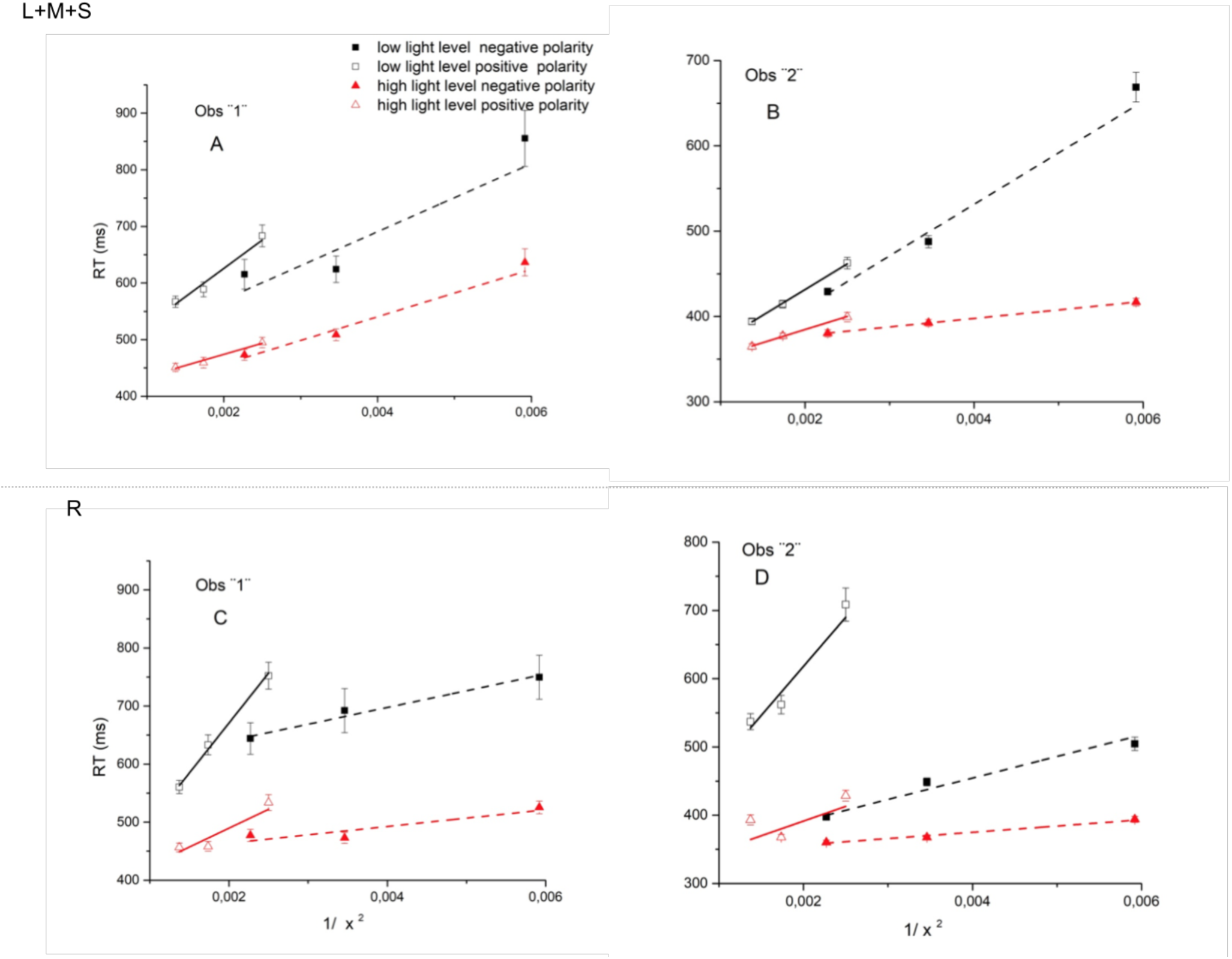
Each panel shows the reaction times as a function of the reciprocal of signal intensity, using the light level and polarity as labeled for both observers 1 and 2. A and B panels correspond to RTs data for only cones combination (L+M+S) and, C and D panels to RTs data for isolated rods (R). The data have been fitted using Equation (2).

For the R condition (Fig. 3, lower panels), no differences were found for the intercept (Obs. 1: *t* = 1.82, *p* = 0.210; Obs. 2: *t* = 0.23, *p* = 0.839) nor for the slopes (Obs. 1: *t* = −2.64, *p* = 0.118; Obs. 2: *t* = −1.10, *p* = 0.386) at high light level. However, for low light level, both observers showed significant differences at the slope (Obs. 1: *t* = −11.13, *p* < 0.01; Obs. 2: *t* = −5.17, *p* < 0.05), but only for observer 1 there were significant differences regarding the intercept (*t* = 9.04, *p* < 0.05), more information can be found in the Supplementary Material.

These results indicated that symmetry was achieved at the low light level for the condition L+M+S and at the high light level for the R condition. In the rest of the conditions, the data is not conclusive.

##### Light level effect

For the L+M+S condition (Fig. 3, upper panels), no differences were found for the intercept at both polarities and for both observers (supplementary material). Instead for slopes, at both polarities, no differences were found for observer 1 (positive: *t* = −4.04, *p* = 0.056; negative: *t* = −1.10, *p* = 0.385), but significant differences were found for observer 2 (positive: *t* = −13.97, *p* < 0.01; negative: *t* = −9.44, *p* < 0.05).

For the R condition (Fig. 3, lower panels), no differences were found between both light levels regarding the slopes (supplementary material). Concerning the intercepts, significant differences were found for negative polarity only for observer 1 (*t* = −4.93, *p* < 0.05), for the rest of the conditions no differences were observed (Supplementary material).

From this isolated data, a light level effect was notorious only for one observer in the L+M+S condition, since the slopes were significantly different between light levels for both polarities. The intercept (RT0) for this condition was similar between light levels, which was expected since it suggests processing by one mechanism. Therefore, for the tested range, higher light levels produced lower reaction times.

#### Interaction analysis (L+M+S+R)

##### Polarity effect

Regarding this interaction condition (Fig. 4), no differences were found between polarities at the high light level for both observers, nor for observer 1 at low light level (Supplementary material). Instead, for the observer 2, significant differences were obtained at the low light level (intercept: *t* =-6.92, *p* < 0.05; slope: *t* = 5.24, *p* < 0.05). Therefore, symmetry was found only at high light level; mimicking the isolated R condition.

**Fig. 4.**
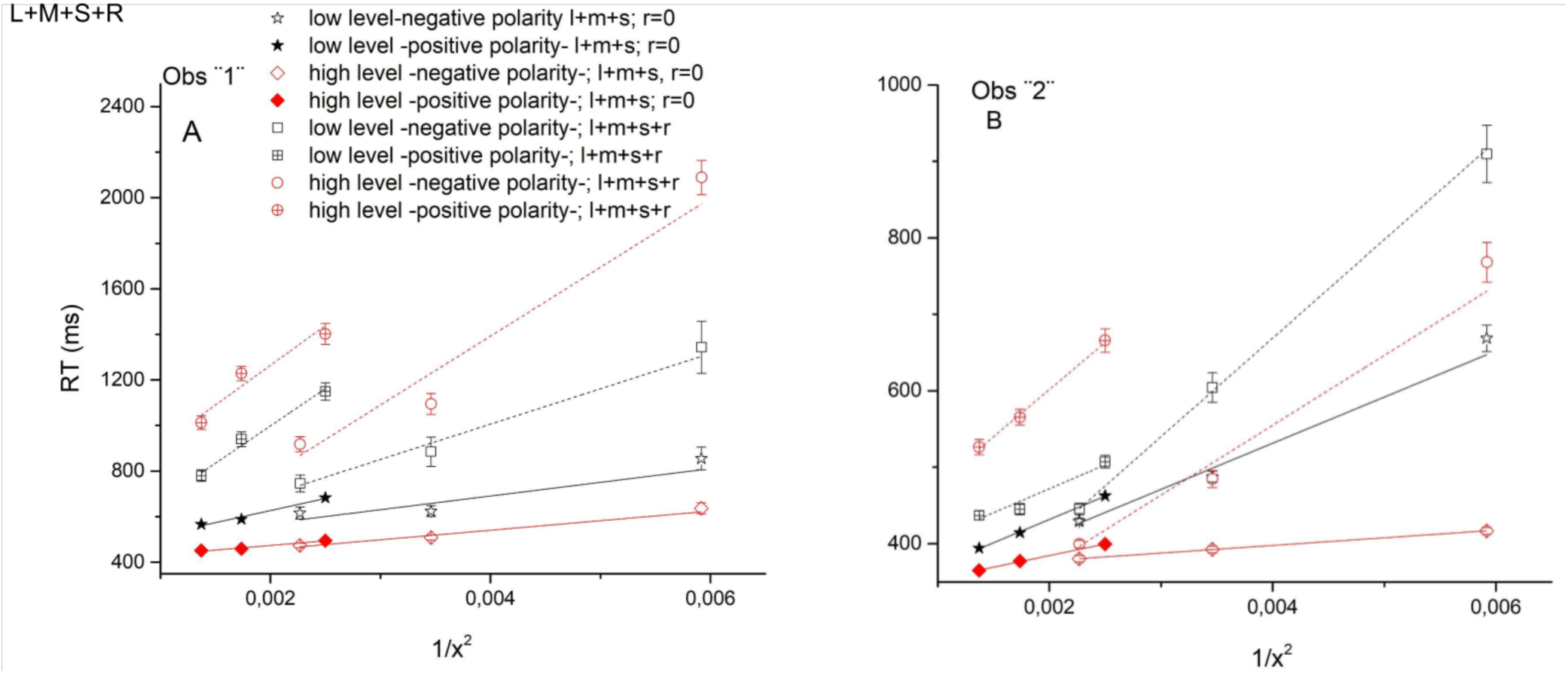
Each panel shows the reaction times as a function the reciprocal of signal intensity, using the light level and polarity as labeled for both observers 1 and 2, including isolated cone contributions (L+M+S; R=0) in top row and interaction between cones and rods (L+M+S+R) in bottom row. The data have been fitted using Equation (2).

##### Light level effect

From this interaction data, we found that although there was a tendency for low light levels to produce lower reaction times in some conditions (Fig. 4), between both light levels no differences were observed at both polarities and for both observers (Supplementary material).

##### Rod intrusion effect

We also compared the L+M+S+R condition with the L+M+S condition in order to assess the rod intrusion effect. From visual inspection of Figure 4, the L+M+S+R RTs seem higher than RTs of L+M+S condition.

Indeed, for both observers, the slopes between both conditions were significantly different in most of the conditions, while the intercepts were mostly similar (Supplementary Material). Therefore, by adding rod stimulation to the luminance pathway, the RT function of the rod-cone interaction was higher than the isolated cones condition RT function (Fig. 4).

As a conclusion of this experiment, we showed that the intrusion of rods in the MC-pathway increased the reaction times, therefore the visual efficiency deteriorated. Furthermore, these results confirmed previous reports studying mesopic reaction times of rods and cones in the MC-pathway (for example [19]), and therefore, this experiment has contributed to validate the methodology used in this work.

### Experiment 2: Intrusion of rods in the PC-inferred pathway

The contributions of rods to the PC-inferred pathway RTs (orange dots) are shown in Fig 5. Also, results for the isolated chromatic pathway (*L/(L+M)*, green dots) and isolated rods (*R*, purple dots) stimulation conditions are shown in the figure. For each receptor type, mean RTs decreased with increments in excitation contrast.

**Figure 5.**
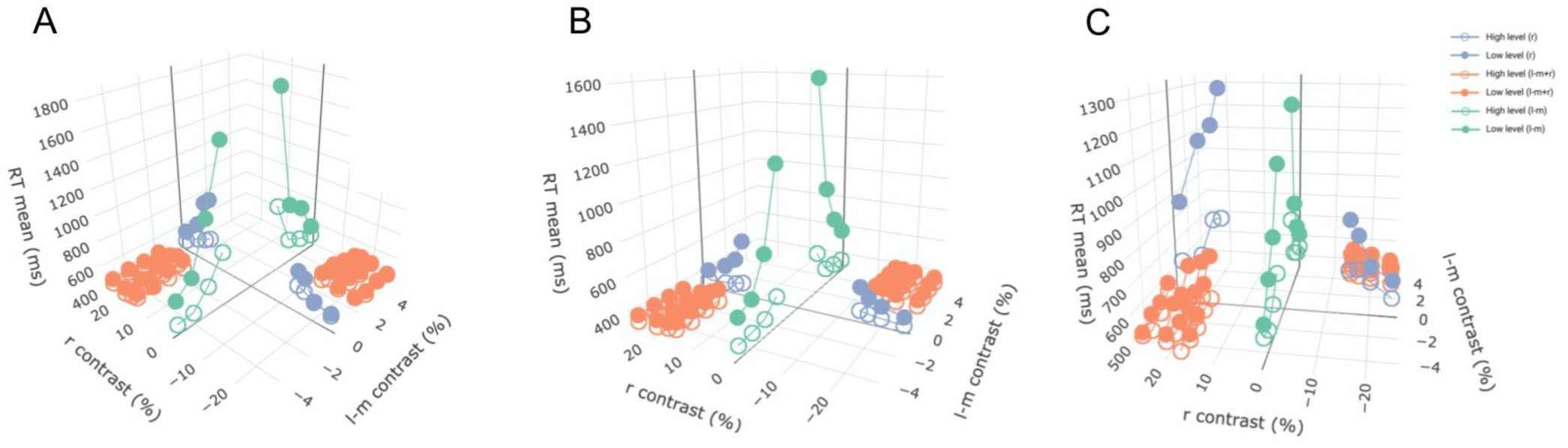
Three-dimensional graph of the results of Experiment 2 for the three observers (A, B, and C). Each panel contains the data of each observer in the Pieron’s space (before linearization). The independent variables of cones L/(L+M) on one axis and rods (*R*) on the other axis. The vertical axis represents the mean of the reaction times in milliseconds [*RT mean (ms)*]. Results are shown for isolated rod condition (purple dots), isolated PC-inferred pathway condition (green dots), and intrusion of rods to PC-inferred pathway condition (orange dots).

In order to better understand the effect on RTs of rod intrusion in the PC-inferred pathway, a thorough analysis was performed (see Methods section). Data (dots) and model results (lines) are shown in Figs. 6 and 7.

**Fig. 6.**
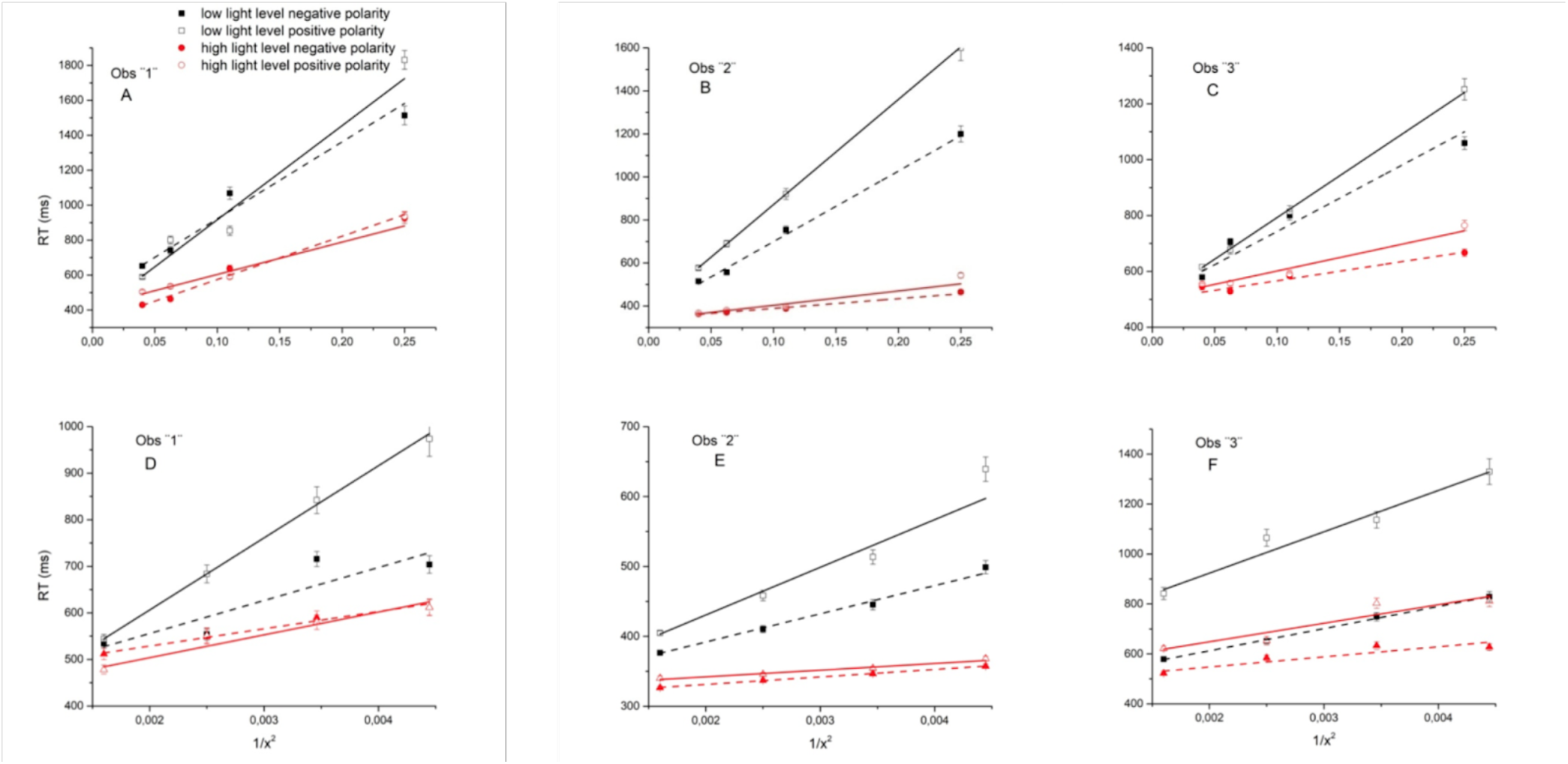
Each panel shows the reaction times as a function of the reciprocal of signal intensity, using the light level and polarity as labeled for three observers, including isolated cone contributions (L/(L+M) in top row and only rods in bottom row. The data have been fitted using Equation (2).

**Fig. 7.**
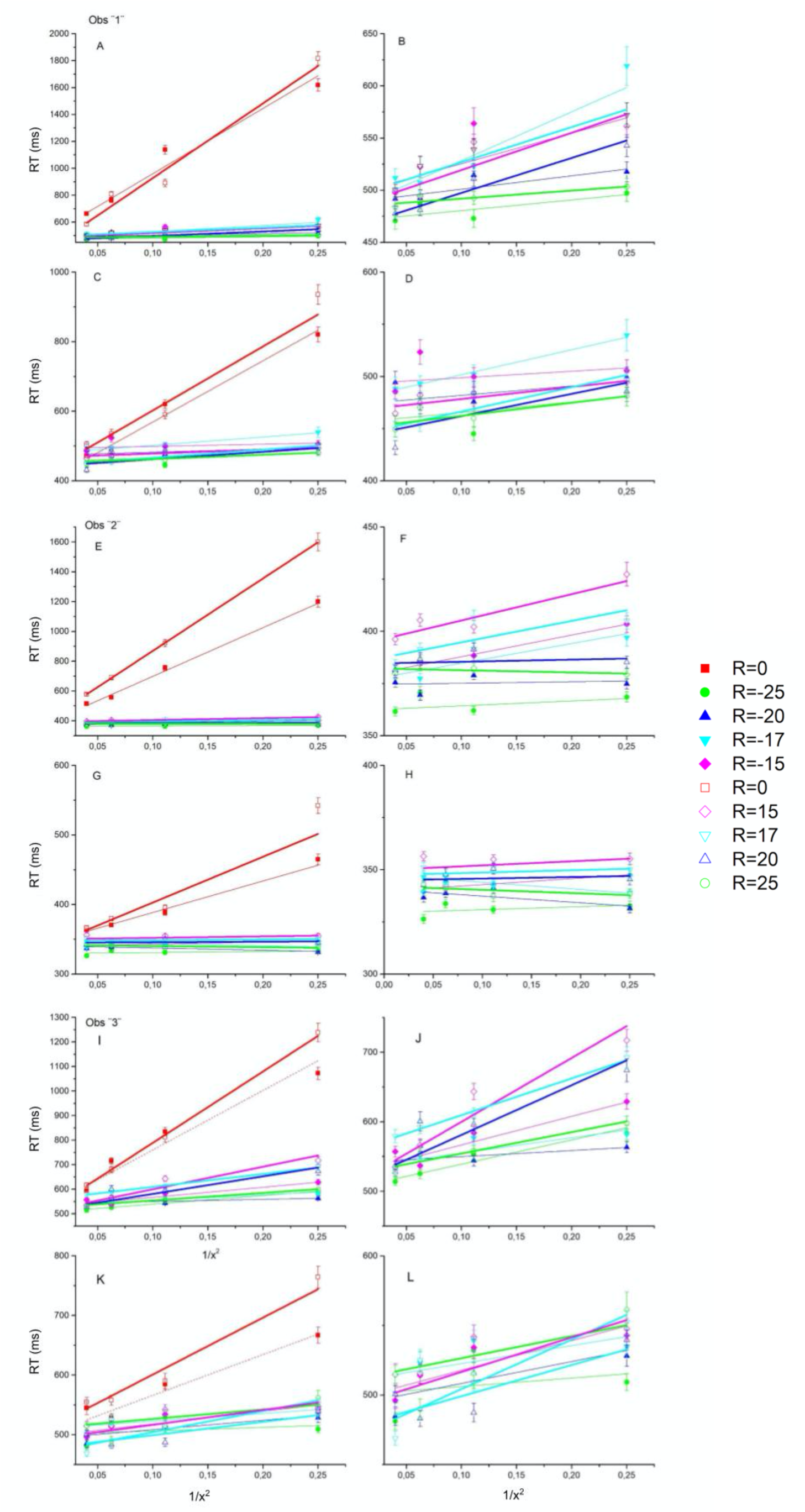
Each panel shows the reaction times as a function of the reciprocal of signal intensity [x = L/(L+M)], using rod value as labeled, for positive and negative contrasts. Both columns contain the RTs for different combinations of L/(L+M) for low light level (panels: A, B, E, F, I, and J) and high light level (panels: C, D, G, H, K, and L). The case where R = 0 contrast (red lines) is only included in the left column. The right column contains same data without R = 0. The data have been fitted using Equation (2)

#### Isolated contributions [L/(L+M), R]

##### Polarity effect

For the *L/(L+M)* stimuli condition, the intercept parameter (RT0) for increments did not have significant differences with respect to those for decrements in high light level and at low light level (Supplementary material). Regarding the slope, there were no differences at both light levels for observers 3 and 1 (Supplementary material), but for observer 2, differences were obtained at both light levels (high: *t* =-3.15, *p* < 0.05; low: *t* =-12.3, *p* < 0.001). It means that at least for two observers, these stimuli conditions were symmetric regarding polarity (see also Supplementary material).

For the R stimuli condition, overall no differences were found regarding polarity (Supplementary material).

##### Light level effect

Comparing results between light levels for the *L/(L+M)* stimuli condition, we observed significant differences in slopes for positive (Obs. 3: *t* = −14.11, *p* < 0.001; Obs. 2: *t* = −36.4, *p* < 0.001; Obs. 1: *t* = −5.6, *p* < 0.01) and negative (Obs. 3: *t* = −4.62, *p* < 0.01; Obs. 2: *t* = −22.1, *p* < 0.001; Obs. 1: *t* = −4.75, *p* < 0.01) contrasts. No differences were found for intercepts at both polarities for observers 3 and 1 (Supplementary material), however for observer 2, a difference was obtained for positive contrasts (positive: *t* =-3.89, *p* < 0.05). It means that RTs of low light level are higher than the RTs of high light level, but both performances tend to have the same value in infinite contrast, at least for two observers.

For R stimuli, differences between light levels were obtained for the three observers, with higher reaction times for low light levels (Supplementary material).

This means that the RTs decreased with increasing luminance levels for both isolated cones and rods as was expected.

#### Interaction analysis [L/(L+M) + R]

The results of the combined *L/(L+M)* condition with different rod contrast are shown in Fig. 7. For the three subjects, and all conditions there is an important overlapping of the interaction data (Fig. 7, right column). There is a tendency to decrease reaction times with increments in absolute rod contrast. However, no significant differences were found when we analyzed both the slopes and the intercepts in most of the interactions regarding polarity and light level (Supplementary material).

#### Rod contribution to the PC-pathway [L/(L+M) vs L/(L+M) + R]

From Fig. 7 (left column), the impact of rod intrusion in the *L/(L+M)* condition becomes evident. In these graphs, the results for the interaction were plotted together with the isolated *L/(L+M)* condition, i.e. no rod intrusion (red lines). When a rod contrast excitation was incorporated, such as values between 15% and 25%, a reduction in reaction time was clearly perceived. This behavior was confirmed statistically. Although the intercepts didn’t differentiate, the slopes were significantly different in almost all cases (Supplementary material), and this difference was more noticed for low light level than for high light level (Fig. 7).

These changes in slopes mean that by adding rod stimulation to the *L/(L+M)* condition, the RTs were faster in the contrast range that we assessed. In other words, the cone-rod interaction improves the RTs for the parvocellular channel under these conditions.

#### Minimum reaction time

Since for each observer significant differences for intercepts were not found among all the conditions tested, we could compute the minimum reaction time predicted by the linear model. The average minimum reaction time was 421 ms for observer 1, 354 ms for observer 2, and 514 ms for observer 3.

## 4. Discussion

We assessed the effect of rod intrusion in reaction times for luminance (MC-inferred) and chromatic (PC-inferred) pathways. The main finding of this study was that rod intrusion reduced the RTs in the PC-inferred pathway, therefore rods improved the performance in vision driven by this chromatic pathway. Interestingly, we found an opposite effect with luminance signals, where rods delayed the responses when they were included in the stimulation of the MC-inferred pathway.

Based on the linearization of the data, we analyzed our results in terms of slope and intercept. For the MC-pathway at low light level, we found symmetry between positive and negative cone (L+M+S) contrasts, which is in agreement with previous psychophysical [10,38,41,42] and physiological [43] reports. The asymmetry found for rods is also supported by the psychophysical literature, Cao and colleagues [40] reported shorter reaction times for negative than positive contrasts which is consistent with our findings (Fig. 6). However, for the light levels in the high mesopic range, our data analysis was not decisive. Previous results of latency studies of ON and OFF pathways are inconclusive, some psychophysical studies suggested that the ON pathway was faster than the OFF pathway in the peripheral retina for both rod and cone systems (e.g. [22]), while others have reached the opposite conclusion [24,25]. Our results support this last conclusion for the rod system.

Our experiments were carried out in two mesopic adaptation levels. Regarding the MC experiment, although for isolated conditions there is a tendency to produce lower reaction times with higher light adaptation levels, this difference was only significant for one observer in one condition. This trend was consistent with previous studies, analyzing the effect of light adaptation in the MC-pathway [12,42]. We couldn’t find clear differences between the results of both light levels for most conditions. Also, for the interaction condition, the apparent flip of the isolated condition trend was not significant. Instead, for the PC pathway, adaptation to higher light levels produced lower reaction times in the isolated conditions, in agreement with previous results [38]. It is interesting to note that the effect of the intrusion of rods in the PC-pathway is stronger for the low light level, in agreement with a higher weight of the rod contribution at this light level [12,44].

For a visual performance task, where two competing independent signals are shown together, one might expect that motor response was driven by the faster one. However, our results showed that an interaction is performed before the decision process to trigger the motor response. This interaction is different with respect to the post-receptoral pathway involved.

For the MC-inferred pathway, our results showed that this interaction is suppressive, in agreement with previous reports assessing the temporal dimension of rod-cone interaction [11,19,23,45–49]. Furthermore, this suppressive interaction could be linear based on previous findings assessing the role of rods activation in the MC-pathway [11,12,18]. From previous findings, this interaction could be at the retinal level [6].

Instead, for the PC-inferred pathway, our results showed a surprising behavior: PC signals become faster when rods are involved. From extra-retinal physiology, it is well known that cells of the PC-pathway have lower temporal sensitivity than cells from the MC-pathway [3,14]. On the other hand, weak rod contributions to the PC-pathway were reported [13,45]. Based on this evidence we might speculate that rods are mainly transmitted via the phasic MC-pathway and the interaction with pure PC-signals is after retinal processing. Furthermore, its nature might follow a probability summation as Sun and colleagues have suggested [11].

From our PC-inferred data, we found no differences in the minimum reaction time, for each observer, among tested stimuli conditions. These RTs range between 354ms and 514ms. This clustering or invariance of RT0 within observers could suggest that there are individual differences but the same process of interaction between rods and cones is verified on every of them. Data from more observers is needed to achieve a stronger conclusion.

The four-primary photostimulator is a tool that made important advances in the study of mesopic vision [11,26]. However, this setup is limited in its spatial possibilities. The arrangement introduced in this study made it possible to generate spatially complex visual stimulation. This is the first experiment assessing mesopic visual performance for spatially complex rod-cone interaction. Traditional rod-cone interaction experiments were carried out with a center field surrounded by a background and most of them have studied lateral or local rod-cone interactions [4,6,20]. We obtained discrimination reaction times, therefore our results have important implications in some applications in the mesopic light range, for example, mesopic visual functions.

Contribution to the three postreceptoral pathways was inferred from natural image statistics [50]. In this work, we focused on the two major pathways, MC and PC. However, rod intrusion in the koniocellular pathway was also identified by physiological studies [51,52]. Still further experiments need to be conducted in order to assess the interaction of rods signals with koniocellular signals in reaction times. Those experiments are beyond the scope of this study.

## Acknowledgments

CONICET PUE 0114 ILAV. Ministerio de modernización de Argentina – Fulbright Commission BEC.AR Program. Agencia I+D+i PICT 2019-03673.

## Notes

### Competing Interest Statement

The authors have declared no competing interest.

