## Supplementary material for "Temporal integration of rod signals in luminance and chromatic pathways"

**Experiment 1: MC-inferred pathway**

*Isolated contributions (L+M+S, R)*

<u>Polarity (Fig. S1)</u>

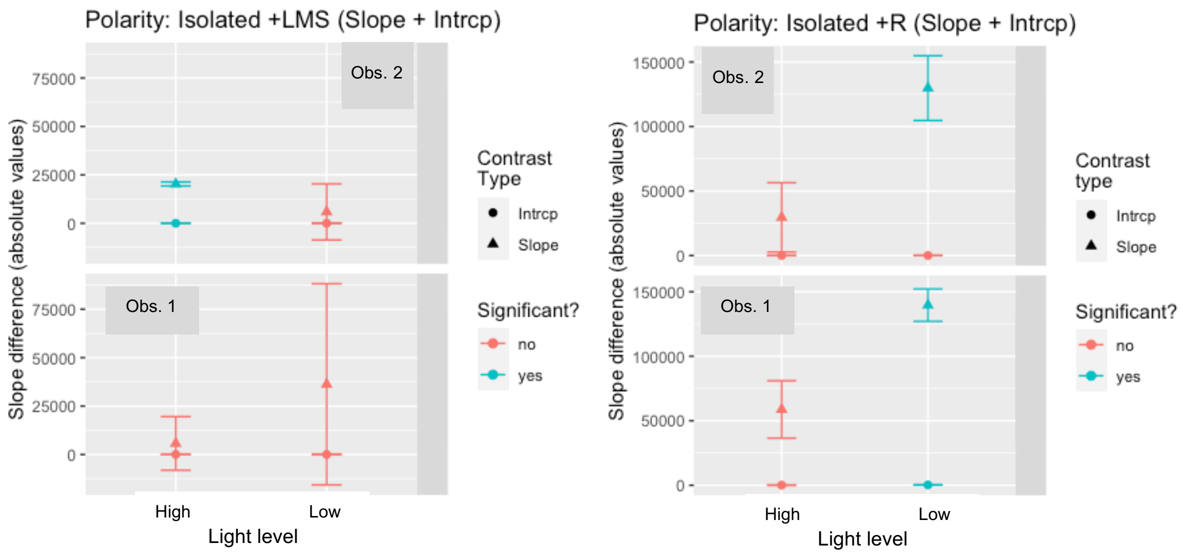


<u>Light level (Fig. S2)</u>

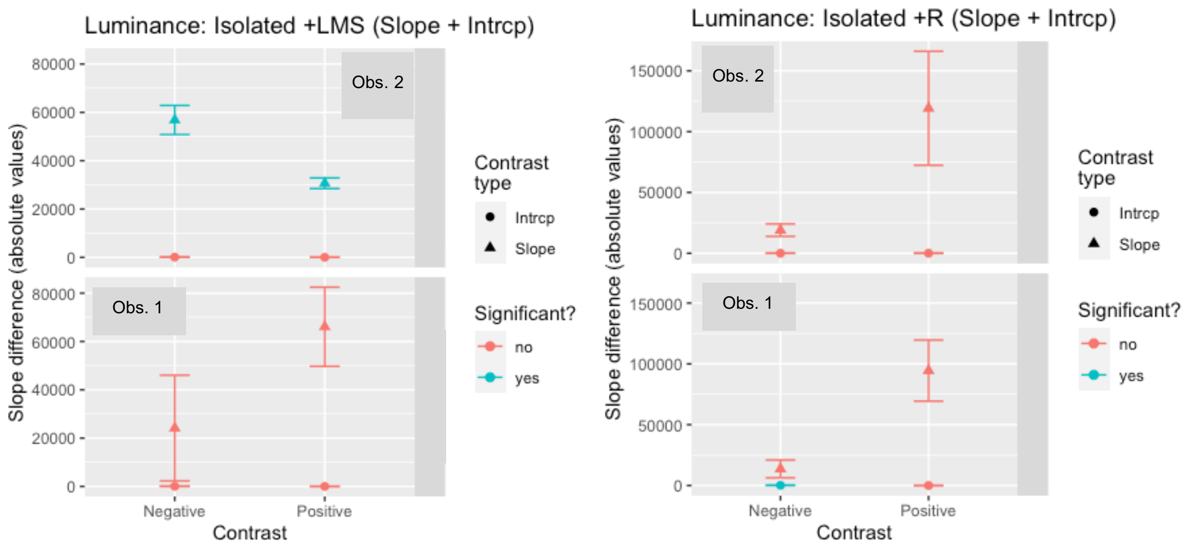

*Interaction contribution (L+M+S+R)*

<u>Polarity and Light level effect (Fig. S3)</u>

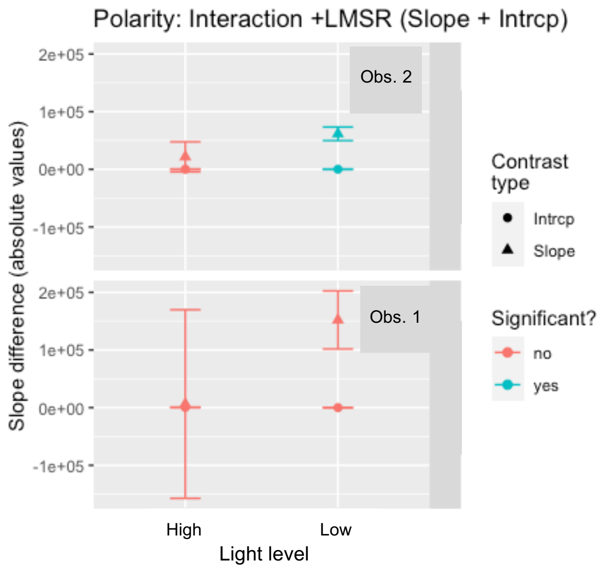


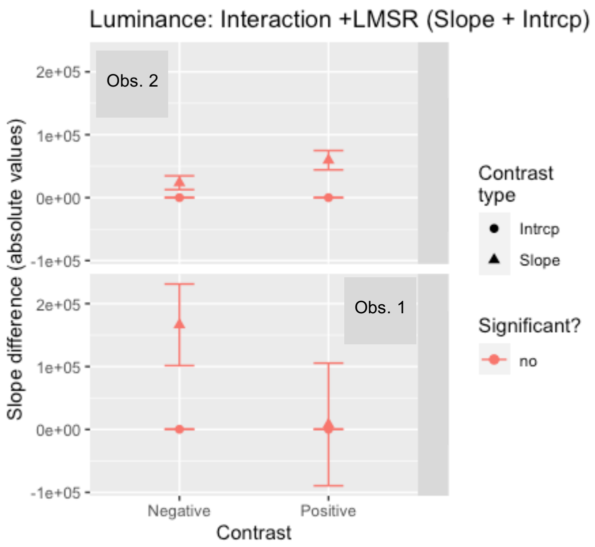


*Rod intrusion effect (Table S1)*

| | | | | Slope | RT0 |
|---|---|---|---|---|---|
| Obs. | Contrast | Polarity | Light level | p-value | p-value |
| 1 | LMS - LMSR | Negative | Low | 0 0.045 | 0.38 |
| 1 | LMS - LMSR | Positive | Low | 0 0.084 | 0.35 |
| 1 | LMS - LMSR | Negative | High | 0 0.070 | 0.54 |
| 1 | LMS - LMSR | Positive | High | 0 0.040 | 0.56 |
| 2 | LMS - LMSR | Negative | Low | 0 0.013 | 0.048 |
| 2 | LMS - LMSR | Positive | Low | 0 0.004 | 0.13 |
| 2 | LMS - LMSR | Negative | High | 0 0.011 | 0.056 |
| 2 | LMS - LMSR | Positive | High | 0 0.800 | 0.37 |

**Experiment 2: PC-inferred pathway**

*Isolated contributions (L/(L+M), R)*

<u>Polarity</u> (Fig. S4)

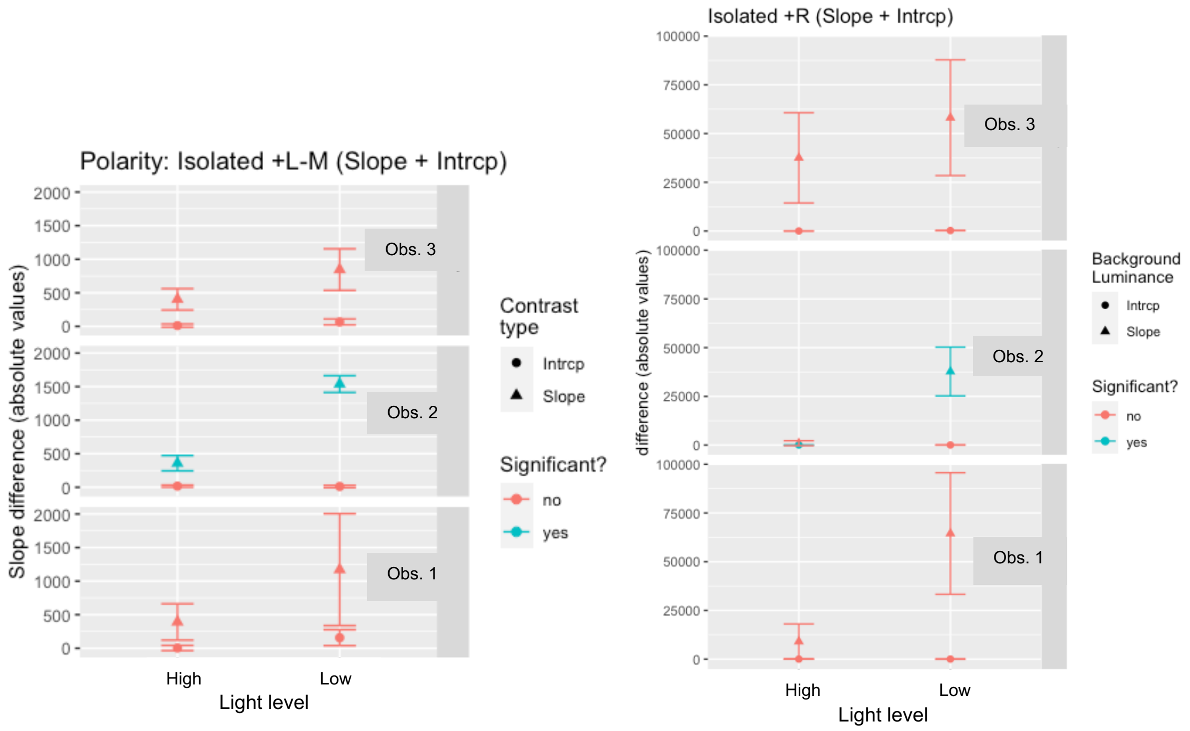


<u>Light level</u> (Fig. S5)

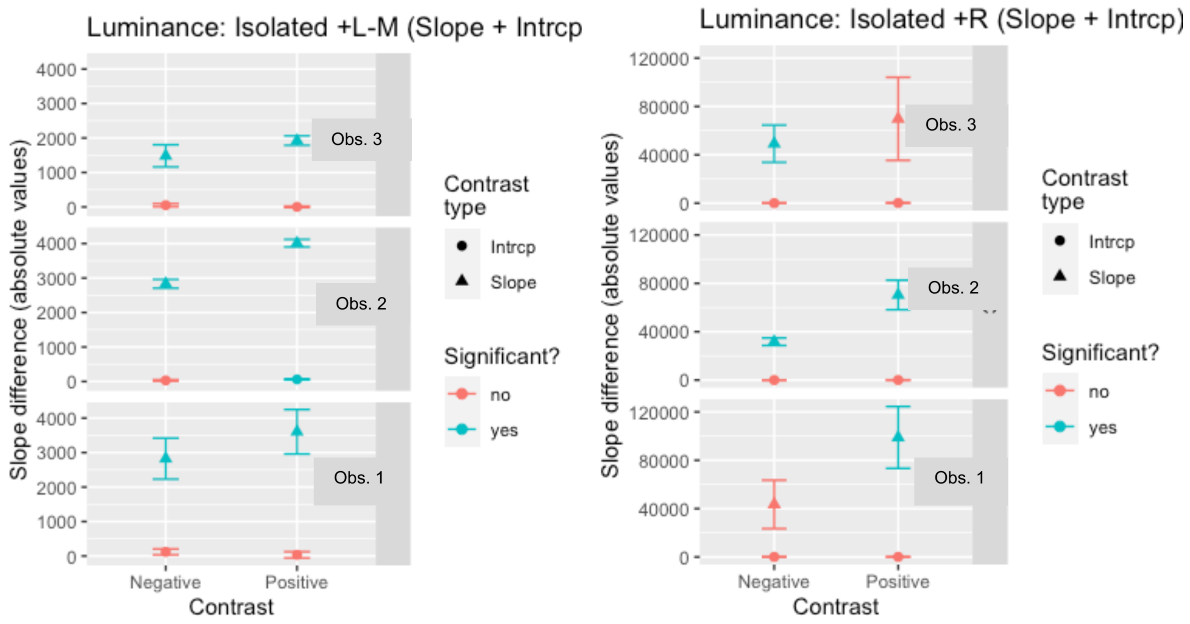

*Interaction analysis [L/(L+M) + R] (Fig. S6)*

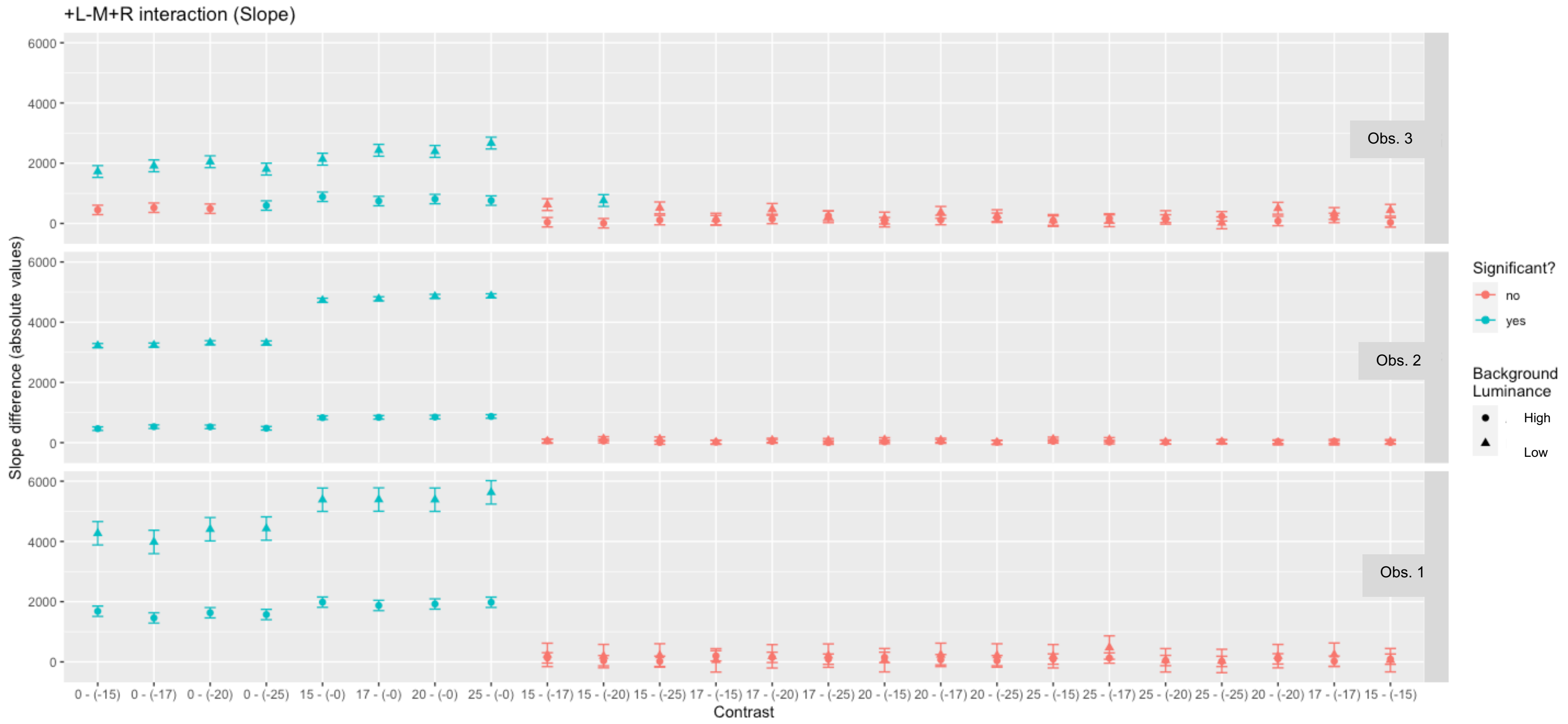

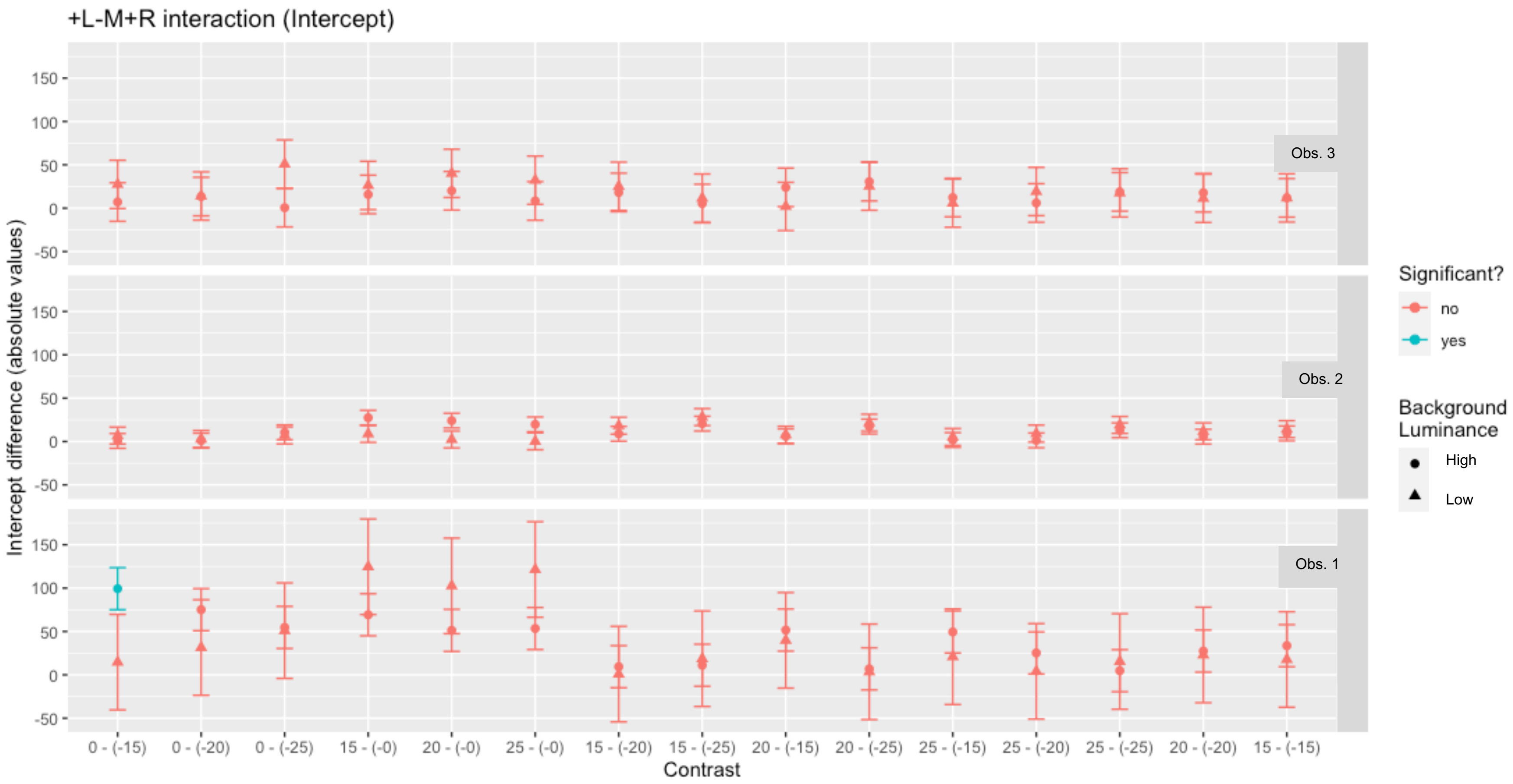
+L-M+R interaction (Intercept)
Intercept difference (absolute values)
Obs. 3
Obs. 2
Obs. 1
150
100
50
0
-50
0 - (-15)
0 - (-20)
0 - (-25)
15 - (-0)
20 - (-0)
25 - (-0)
15 - (-20)
15 - (-25)
20 - (-15)
20 - (-25)
25 - (-15)
25 - (-20)
25 - (-25)
20 - (-20)
15 - (-15)
Contrast
Significant?
no
yes
Background Luminance
High
Low
